# Influence of the large–Z effect during contact between butterfly sister species

**DOI:** 10.1101/2020.06.04.134494

**Authors:** Erik D. Nelson, Qian Cong, Nick V. Grishin

## Abstract

Comparisons of genomes from recently diverged butterfly populations along a suture zone in central Texas have revealed high levels of divergence on the Z chromosome relative to autosomes, as measured by fixation index, *F*_*st*_. The pattern of divergence appears to result from accumulation of incompatible alleles, obstructing introgression on the Z chromosome in hybrids. However, it is unknown whether this mechanism is sufficient to explain the data. Here, we simulate the effects of hybrid incompatibility on interbreeding butterfly populations using a model in which populations accumulate cross–incompatible alleles in allopatry prior to contact. We compute statistics for introgression and population divergence during contact between model butterfly populations and compare them to statistics obtained for 15 pairs of butterfly species interbreeding along the Texas suture zone. For populations that have evolved sufficiently in allopatry, the model exhibits high levels of divergence on the Z chromosome relative to autosomes in populations inter-breeding on time scales comparable to periods of interglacial contact between butterfly populations in central Texas. Levels of divergence on the Z chromosome increase when interacting groups of genes are closely linked, consistent with interacting clusters of functionally related genes in butterfly genomes. Results for various periods in allopatry are in qualitative agreement with the pattern of data for butterflies, supporting a picture of speciation in which populations are subjected to cycles of divergence in glacial isolation, and partial fusion during interglacial contact.

## Introduction

Recent studies comparing divergent populations of butterflies have revealed elevated levels of divergence on the Z chromosome relative to autosomes [1, 2]. To explain these observations, it was suggested that the observed patterns of divergence result from the accumulation of postzygotic incompatibilities, obstructing introgression on the Z chromosome in hybrids (see, for example, Fig. 5 of reference [2]). However, it is known that a number of other factors can contribute to this effect, such as changes in population size, differing rates of reproductive success for male and female butterflies, and a smaller effective population size for Z chromosomes relative to autosomes [3]. As a result, it is of interest to know the “bare” contribution of hybrid incompatibilities to the extent of divergence in autosomes and Z chromosomes, e.g. in a model where these effects are absent and mutations are individually neutral. In this work, we develop such a model and compare our results to those obtained by Cong et al. [2] for 15 closely related species of butterfly interbreeding along a suture zone in central Texas.

The central Texas suture zone is formed by emigration of species from glacial refugia along coastal and inland regions of Mexico and the southern United States, including the Yucatan peninsula and the state of Florida (see Fig.s S1 and S2). The species sampled by Cong et al. diverged on the order of 1 million years ago [4], and have, as a result, experienced multiple periods of glacial cooling and interglacial warming. During glacial periods, central Texas was subjected to severe decreases in temperature [5], which would have caused drastic, if not total isolation of sister species in southeastern and southwestern refugia; During the most recent warming period, sister species migrated into Texas, while major portions of the populations remained in refugial regions, isolated from the suture zone by large distances. To determine the influence of hybrid incompatibilities with the Z chromosome during contact, we will at first neglect the effects of isolation by distance, and consider a generic model of secondary contact [6, 7] in which a population divides, the resulting sister populations evolve for a period in allopatry while accumulating hybrid incompatibilities, and later begin to interbreed. We then compare measures of introgression and population divergence for gene sequences in our model during periods of contact to results obtained for *∼* 1 kb sequence windows by Cong et al. [2].

To represent the state space for pairs of sister populations, Cong et al. employed two basic statistics: The index of gene flow, *I*_*gf*_, a simple extension of the indicator function *G*_*min*_ developed by Geneva et al. [6], defined as the fraction of sequence windows with *G*_*min*_ *≤* 0.25, and the fixation index, or relative divergence, *F*_*st*_. *I*_*gf*_ describes the fraction of sequence windows where introgression has occurred, while *F*_*st*_ describes the degree of genomic difference between populations (we note that *I*_*gf*_ is not directly related to gene flow as measured by the effective migration rate [8]). Multiple genomic samples were collected from each pair of populations, and separate indices were computed for autosomes and Z chromosomes. The results are shown in Fig. 1; The data points in this figure describe index values for sister organisms that have been classified as different species in the literature (green), more closely related organisms for which classification is uncertain (yellow), and samples of the same species (red). When populations are compared through their autosomes (Fig. 1A), the data exhibit a continuous pattern across the entire range of index values; However, for the Z chromosome (Fig. 1B), the data obtained from samples of the same species (red) are separated from those of closely related species by a gap of “missing” values, which could suggest a rapid transition [1,9]. For different species (green and yellow data points), *F*_*st*_ values for the Z chromosome are always larger than those for autosomes (Fig. 2). At the same time, the fraction of divergent nucleotide positions in the Z chromosomes of sister species is slightly smaller than that for autosomes [2], indicating similar rates of adaptation. In accord with these results, Cong et al. have argued that the pattern of data in Fig. 1 reflects the influence of negative interactions between autosomes and Z chromosomes in hybrids during periods of interbreeding – i.e., the large–Z effect [3]. Our goal in this work is to determine whether this mechanism is sufficient to explain the data for butterflies – in particular, the large gaps, Δ*F* = *F*_*Z*_ *− F*_*A*_, between *F*_*st*_ values for autosomes and Z chromosomes shown in Fig. 2. To accomplish this, we simulate populations with different divergence times in allopatry, different migration rates, and different levels of hybrid incompatibility, leading either to fusion or continued divergence during secondary contact. Mutation rates, and rates of recombination within and between gene sequences are varied about values for Drosophila and Heliconius butterflies. As Bank et al. have pointed out [10], negative interactions between individually neutral mutations (i.e., neutral incompatibilities) are unstable in interbreeding populations, and ultimately break down due to recombination [11]. However, when crossovers between genes are infrequent, consistent with closely linked genes on butterfly chromosomes, there is an initial period during secondary contact where Δ*F* can increase dramatically, depending the migration rate and the strengths of interactions between incompatible alleles. In this case, which would perhaps correspond to interacting clusters of functionally related genes [12,13], large values of Δ*F* can develop in fusing populations on time scales comparable to interglacial warming periods; During these periods, the gap of missing *F*_*st*_ values obtained in Fig. 1B is statistically likely over a wide range of interaction strengths and migration rates for populations that have diverged sufficiently in allopatry. Mean values of *I*_*gf*_ during allopatry and secondary contact are in good agreement with Fig. 1 for autosomes, but slightly larger than expected for the Z chromosome. We return to these points later below.

**Fig. 1.**
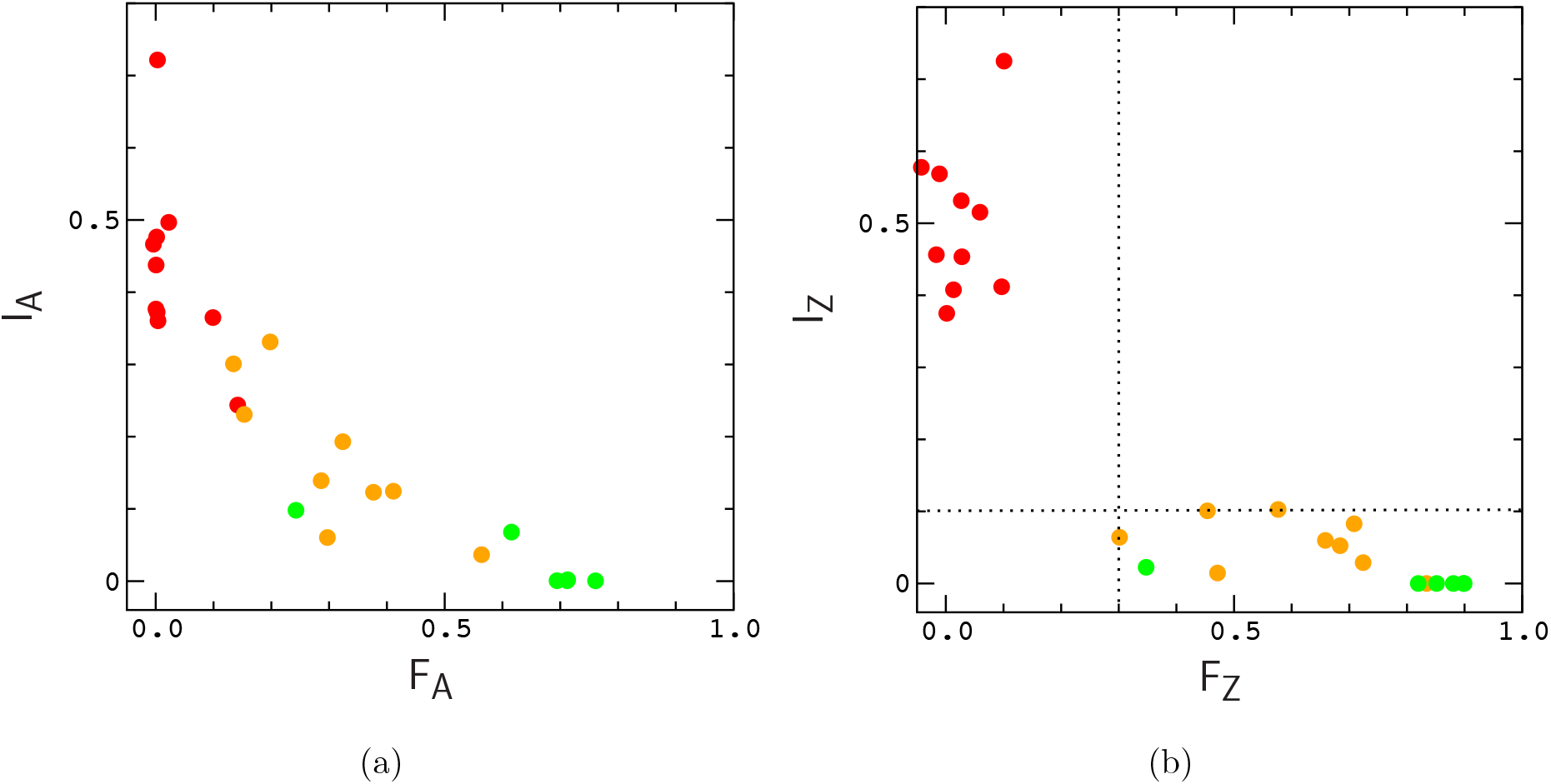
Index of gene flow versus index of fixation for autosomes (A) and Z chromosomes (B) of sister species sampled by Cong et al. Data for *I*_*A*_ and *I*_*Z*_ is multiplied by a factor of 4 to remove a scaling factor used in their work. Data points describe pairs of organisms that have been classified as different species in the literature (green), closely related organisms for which classification is uncertain (yellow), and organisms of the same species (red). The dotted lines in panel (B) are included simply to guide the eye.

**Fig. 2.**
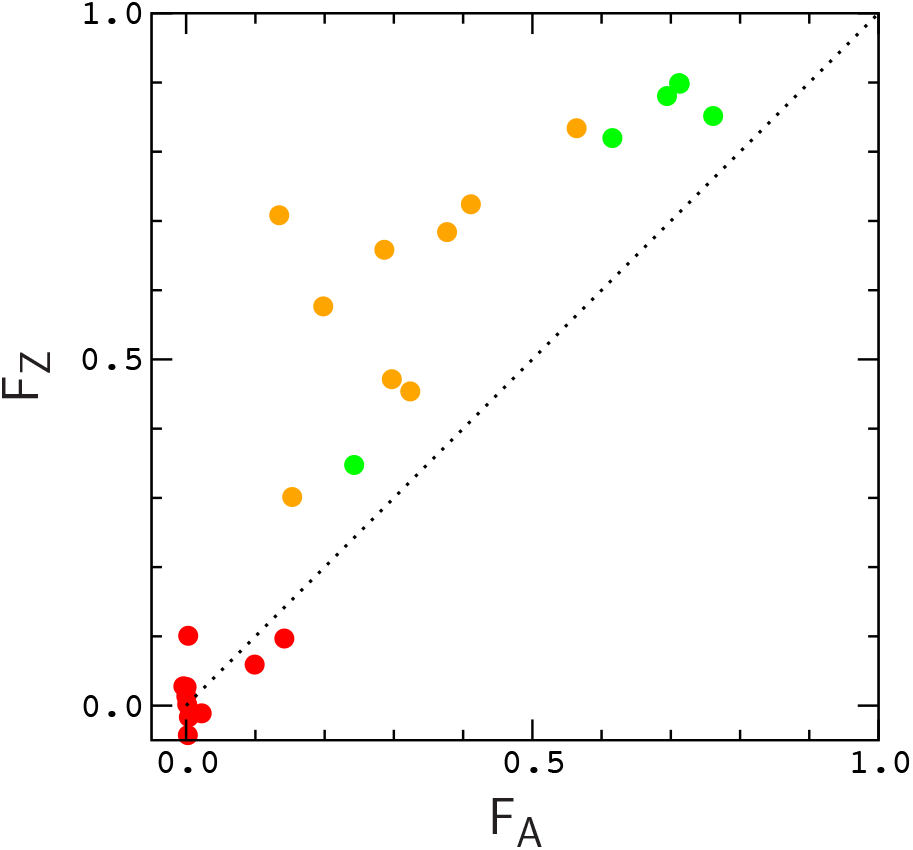
Correlation between *F*_*Z*_ and *F*_*A*_ from Fig. 1. The dotted line *F*_*Z*_ = *F*_*A*_ is included to guide the eye.

## Methods

We simulate populations of diploid individuals evolving in allopatry and secondary contact, and we compute statistics for gene segments, consistent with the approach used in Fig. 1. Model genomes consist of several pairs of chromosomes, each containing a number of binary genes of length *L* loci. Populations evolve by plain Wright–Fisher dynamics with random mating between male and female individuals [14]. In each generation, mutations occur within genes at a rate *µ* per gene per generation. Pairs of male and female individuals are then selected at random according to fitness for mating. Male genomes undergo explicit meiosis, in which chromosomes are duplicated, and the resultant chromatids undergo random crossing over [15], with separate rates, *r* and *r*^*′*^, for crossing over within and between gene segments (meiosis is achiasmatic in model females, consistent with butterfly reproduction [16]). A single offspring is generated from each mating event by random union of male and female gametes, and the process is continued until the original population is replenished. Finally, an equal number of offspring (with mean *Nϵ*, where *N* is the size of a population and *ϵ* is the migration rate) are randomly selected from each population to undergo migration, and the selected individuals are then exchanged between populations (in allopatry, *ϵ* = 0).

We consider two different scenarios for secondary contact: In scenario (i), a population of size *N* is first equilibrated for a period of Δ*t*_*E*_ generations. Let 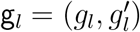 denote the allelic state of a diploid locus *l* in a genome g. Initially, all genomes in the population have g_*l*_ = 0 uniformly, and mutation events act to assign the maternal (*g*_*l*_) or paternal 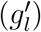 states of a locus to 1. All mutations are individually neutral. After equilibration, loci that have fixed in the population for the mutant allele type are returned to their initial states. The population is then duplicated, and the resultant “sister” populations evolve in allopatry for a period Δ*t*_*A*_. At the end of this period, loci that have fixed for the mutant allele across both populations are returned to their initial state. Several pairs of loci are then selected to participate in hybrid fitness interactions (see below), and the two populations evolve in contact for a period Δ*t*_*C*_ subject to fitness costs incurred due to interactions formed by various allele combinations at the selected loci. In scenario (ii), the entire procedure is the same, except that the initial population has size 2*N* before dividing into two populations of size *N*. In both scenarios, the W chromosome acts only to determine the sex of an individual.

During allopatry, mutant alleles are lost, or rise toward fixation in each population via genetic drift. Loci that are nearly fixed for the mutant allele type in one population are usually far from fixation in the other. Let *p*_*l,γ*_ denote the frequency of the mutant allele type at locus *l* in population *γ*, with *γ* = 1, 2. To the describe the cost of hybridization, we select loci in autosomes for which *p*_*i*,1_ *∼* 1 and *p*_*i*,2_ *∼* 0 to interact negatively with loci in the Z chromosome(s) for which *p*_*j*,1_ *∼* 0 and *p*_*j*,2_ *∼* 1 (see Fig. 3). We then repeat this process with the population subscripts interchanged, selecting an equal number of loci in autosomes with *p*_*i*,2_ *∼* 1 and *p*_*i*,1_ *∼* 0 to interact negatively with loci in the Z chromosome(s) for which *p*_*j*,2_ *∼* 0 and *p*_*j*,1_ *∼* 1.

**Fig. 3.**
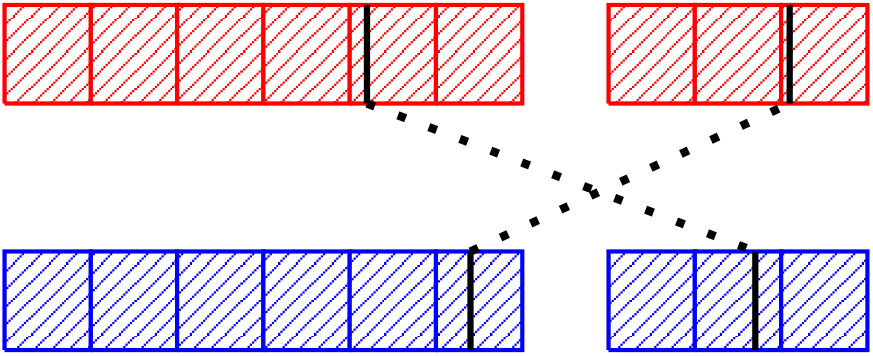
Part of a hybrid genome formed at the time of contact. The illustration describes a pair of hybrid-in3teractions (dotted lines) connecting selected alleles (solid lines) in the Z chromosome (left) to0selected alleles in an autosome (right) in a male hybrid genome. Gene segments are denoted by shaded squares.

Let *f* (g_*i*_, g_*j*_) denote the log fitness cost for a pair of selected loci, (*i, j*). We define *f* as follows: If both loci are homozygous for the mutant allele, *f* = 4*s*; if one locus is homozygous for the mutant allele and one locus is heterozygous, *f* = 2*s*; and if both loci are heterozygous, *f* = *s/*4. For all other combinations, *f* = 0. The fitness of a genome g is then defined as, ω = exp − ∑_(*i,j*)_ *f* (*g*_*i*_, *g*_*j*_). Here, mutant loci on the single Z chromosome of a female genome are dominant, and act as homozygous loci on the two Z chromosomes of a male genome. This condition, and the fact that *f* is less than *s* when both loci are heterozygous, ensures that hybrid females are typically less fit than hybrid males, consistent with Haldane’s rule. As result, gene flow on the Z chromosome is limited by the fitness of female genomes, in accord with the analysis of Cong et al. [2]. For male genomes, the fitness model is basically the same as the “pathway” model used by Lindtke and Buerkle to describe hybrid interactions among autosomal loci [11].

Below, we simulate populations of genomes with three pairs of chromosomes in which autosomes carry three genes, and Z chromosome carry six genes. To define *ω*, we select six pairs of loci for which the differences between *p*_*l*,1_ and *p*_*l*,2_ above are largest. In most of the simulations, we select loci that connect the first pair of autosomes to the Z chromosome(s). The parameters of the simulations are selected so that their scaled values (*Nµ, Nr*, and *Nr*^*′*^) agree in order of magnitude with values obtained for Heliconius and Drosophila, an organism commonly used to infer the biochemical functions of butterfly genes [2]. The scaled mutation rate for a gene (of typical length 1770 bp) in Drosophila is about *Nµ ∼* 1 [17]; Here, we will assume a similar rate for butterflies. The typical length of a chromosome in Heliconius is about 20 Mbp, and the crossover rate per chromosome is about *r ∼* 1 per generation [16]; If we assume that genes in Heliconius are comparable in length to those in Drosophila, we obtain a crossover rate per gene in Heliconius of about *r ∼* 10^*−*4^ per generation. The effective population size for Heliconius is about 10^6^, which leads to a scaled rate of about *Nr ∼* 100 per gene per generation [3].

In practice, we describe genes using strings of characters in our C++ code. The number of mutant alleles participating in a gene segment is typically less than a few percent for the timescales considered here, so that most of the loci in a gene do not carry divergent mutations (see [2] for comparison with butterfly populations). For this reason, we use smaller gene strings of length *L* = 100 loci in our code to reduce the cost of the simulations. We simulate populations of *N* = 10^4^ individuals, while varying the parameters *s*,**ϵ**and *r*^*′*^. Since the morphologies (i.e., sex organs, wing color patterns, etc.) of butterfly sister specimens are very similar, prezygotic barriers to introgression may be small. For this reason, we sample moderate to large migration rates, 0.1 *≤ N*ϵ ≤ 10. To counterbalance the effects of migration, we sample a broad range of interaction strengths, 0.01 *≤ s ≤* 0.1, including the “null model”, *s* = 0, for comparison. Our main findings are summarized in the figures below. Supporting figures are contained in the supplementary material file. Data and C++ code used to conduct the simulations can be obtained from: https://cloud.biohpc.swmed.edu/index.php/s/D766rG6fWZWrsfG.

## Results

To begin our investigation, we first explore the time dependence of the statistics *G*_*min*_ and *F*_*st*_ for populations evolving in allopatry for comparison with the results of Geneva et al. [6]. To define the statistics, let 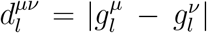 denote the difference (Hamming distance) between genomes g^*µ*^ and g^*v*^ at (haploid) locus *l*, and let

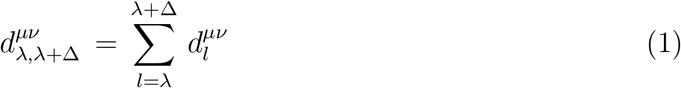

denote the distance between g^*µ*^ and g^*v*^ for a window of loci, [*λ, λ* + Δ]. Assume that we have sampled a small number of genomes from each population. For a given window of loci, *G*_*min*_ is then defined as the ratio [6],

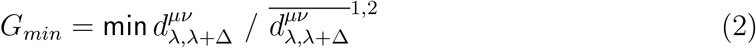

where min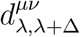 are 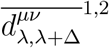 the minimum and average distances between sequencesand sampled from different populations; The fixation index, or relative divergence is defined as [6],

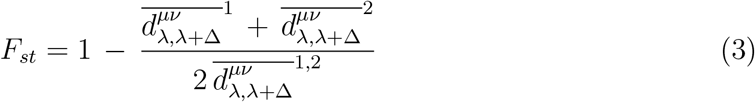

where, for example, 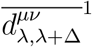 is the average distance between sequences sampled from population 1 with *µ ≠ v*. Below, we compute *G*_*min*_ for individual gene sequences, and we compute *F*_*st*_ by averaging the numerator and denominator of the fraction in Eq. (3) over gene sequences [18]. Unless otherwise noted, we compute *G*_*min*_ by sampling four genomes from each population, and we compute *F*_*st*_ by sampling ten genomes from each population (this is intended as a means to reduce noise in plots like Fig. 1). The introgressionmeasure, *I*_*gf*_, is defined as the fraction of gene segments with *G*_*min*_ *≤* 0.25 (see Fig. 4Aof Geneva et al. [6] for comparison). Except for the number of samples used to compute *F*_*st*_, our approach is the same as that used by Cong et al. [2].

**Fig. 4.**
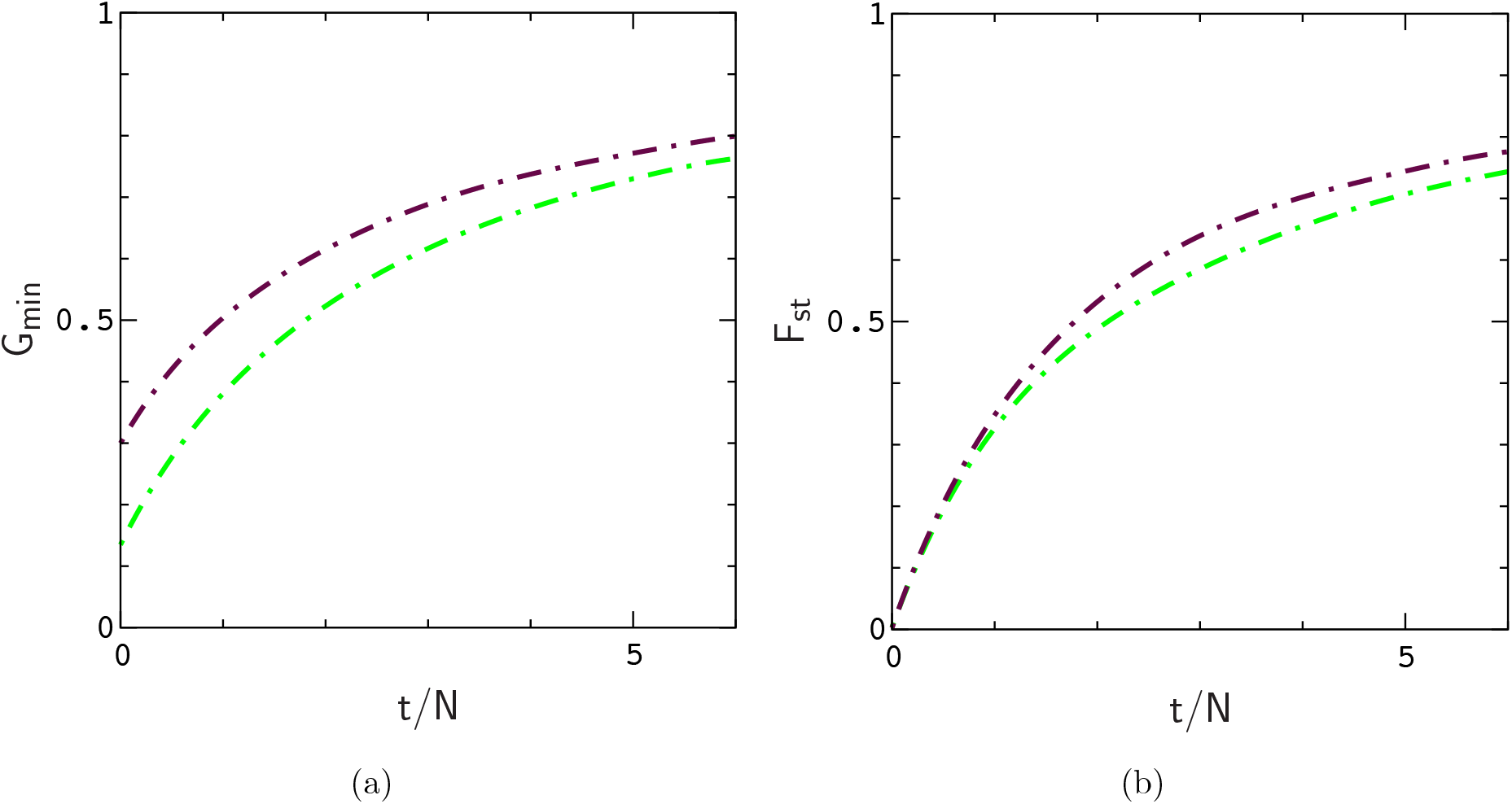
Mean value of *G*_*min*_ and *F*_*st*_ for autosomal genes as a function time since diverging in allopatry under scenarios (i) (green) and (ii) (maroon). Averages are computed from 128 replicate simulations with *N* = 10^4^, *µ* = 10^*−*4^, and *r, r*^*′*^ = 10^*−*2^. The plots are precise polynomial fits to the averages.

**Fig. 5.**
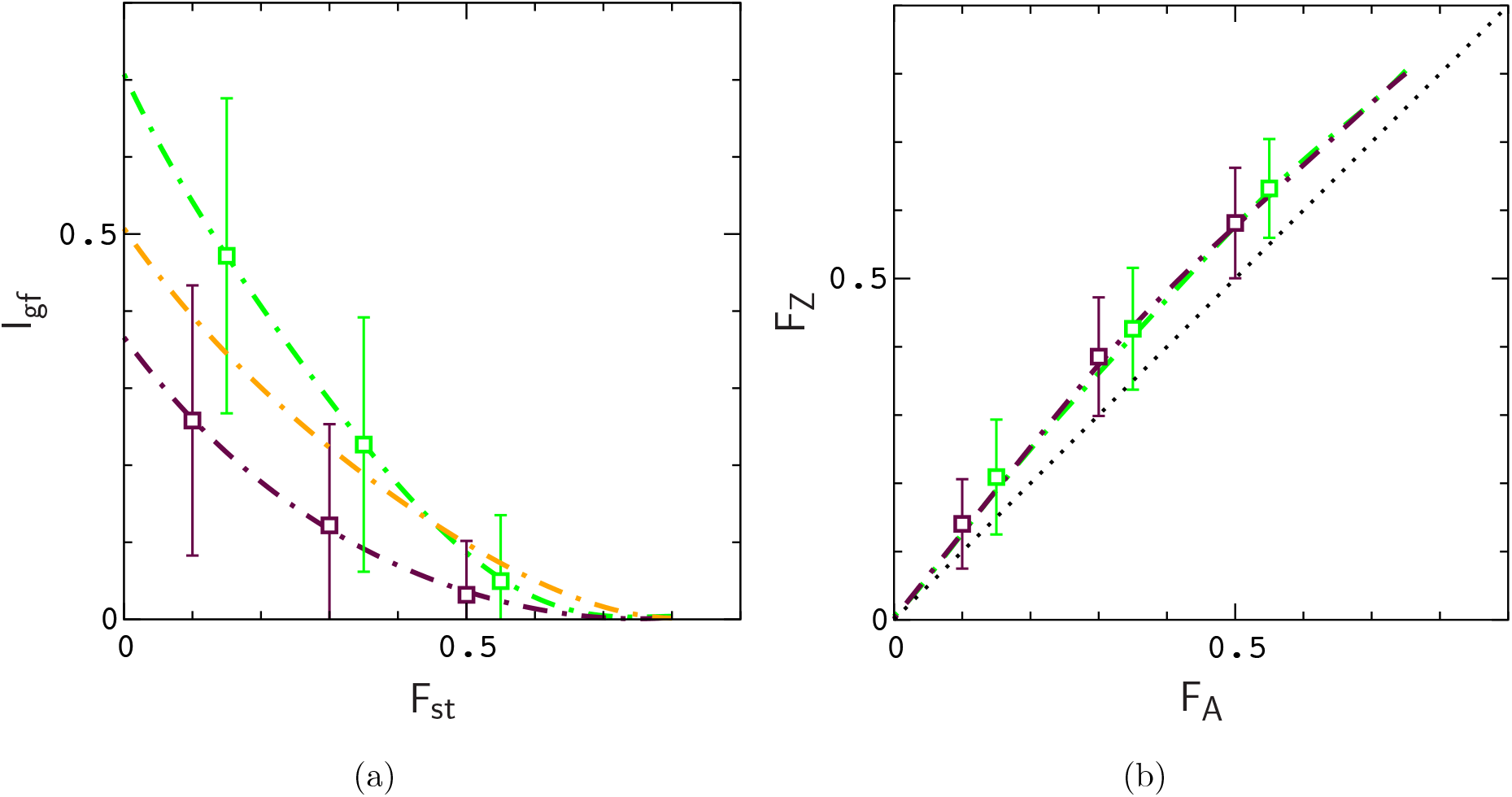
Mean value of *I*_*A*_ and *F*_*Z*_ versus *F*_*A*_ (squares) corresponding to the simulations in Fig. 4. Error bars denote the widths of the distributions. Broken lines are precise polynomial fits to the paths defined by ⟨*I*_*A*_⟩ (*t*) and ⟨*F*_*Z*_⟩ (*t*) versus ⟨*F*_*A*_⟩ (*t*) as a function of time, where braces denote averaging over simulations (a plot of ⟨*I*_*Z*_⟩ (*t*) versus ⟨*F*_*Z*_⟩ (*t*) for scenario (ii) (orange) is also included for comparison with Fig. 1). The widths of the distributions for *I*_*gf*_ reflect the small number of gene sequences considered in the model.

In Fig. 4, we plot *F*_*st*_ and *G*_*min*_ for autosomal genes as a function time since diverging in allopatry under scenarios (i) and (ii). The results can be compared with those in Fig. 2 of Geneva et al. [6]. Although a direct comparison is not possible (Geneva et al. average simulations of a single sequence window over a range of *µ* and *r* values), our results behave as one would expect for the lower values of *µ* and *r* used in our simulations (note that the transition to allopatry in Geneva et al. is analogous to our scenario (i)). Interestingly, there is a noticeable difference in the plots of *G*_*min*_ for duplication and division of populations in allopatry, and we find that scenario (ii) leads to closer agreement with butterfly data for *I*_*gf*_ (Fig. 5). Plots of *F*_*st*_ computed from four and ten samples per population are essentially identical (this is not surprising since *F*_*st*_ already reflects an average over several gene windows).

In the remaining figures, we describe results for secondary contact between populations under scenario (ii). To plan our simulations, we assume that periods of contact are comparable to those of real populations during interglacial warming periods. The time scale for interglacial periods in North America over the last million years is roughly between 10^4^ and 10^5^ years. To calibrate the model to real time scales, we assume, consistent with our choice of parameters, that *N* generations in the model correspond to *N*_*e*_ generations for butterfly populations, where *N*_*e*_ is the effective population size for butterflies. Then, solving for *α* in the expression *α N*_*e*_*τ* = Δ*τ*, where *τ* is the generation time and Δ*τ* is the length of an interglacial period, the corresponding period of contact in the model is *αN*. Although data for *N*_*e*_ is unavailable for the species in Fig. 1, we can obtain a rough idea of how *N*_*e*_ varies over time and among species from the study of Heliconius populations by van Belleghem et al. [3] (for Drosophila, see [19]). Below, we focus our attention on values of *N*_*e*_*τ* on the order of 10^5^ years, consistent with the lower range of *N*_*e*_ values in van Belleghem et al. [3], in which case, *N* generations in the model corresponds to the length of a glacial or interglacial period for butterflies. We then simulate different values of ϵ and *s* for different periods in allopatry.

The results of this survey are summarized in Fig.s 6–9. In Fig.s 6 and 7 we plot the mean value of *F*_*Z*_ versus *F*_*A*_ during contact for populations that have evolved in allopatry for Δ*t*_*A*_ = *N* and Δ*t*_*A*_ = 2*N* generations; The mean values of *F*_*A*_ just prior to contact are *F*_*A*_ 0.3 and *F*_*A*_ 0.5, respectively. Populations remain in contact for Δ*t*_*C*_ = *N* generations. Each set of averages (circles, squares, etc.) is the result of 128 replicate simulations (i.e., with the same values of *s* andϵ) sampled every 100 generations. Lower panels in the figures describe the fraction of simulations contributing to each data point The numbers of samples per data point are shown in Fig.s S3 and S4. Of particular interest is the index value *F*_*A*_ 0.15, the smallest value of *F*_*A*_ for different species (yellow and green data points) in Fig. 2. As is evident by inspection of the data for *F*_*A*_ 0.15 in Fig. 6B, large values of Δ*F*, consistent with the data for butterflies (Fig. 2), can occur at low frequency if hybrid interactions are sufficiently strong. For weaker interactions (Fig. 7), such that small values of *F*_*A*_ are frequent, the mean value of *F*_*Z*_ still usually remains above the point *F*_*Z*_ 0.3 corresponding to the lower edge of the data for different species in Fig. 1. As a result, samples taken from a simulation during contact are unlikely to occur in the region of missing *F*_*Z*_ values. A similar pattern emerges for shorter times in allopatry, with more weakly diverged populations (Fig. 6A); Because fewer loci reach fixation when Δ*t*_*A*_ is small, stronger interactions are required to maintain a significant level of divergence for the Z chromosome. In all of the simulations, mean values for *I*_*A*_ versus *F*_*A*_, and *I*_*Z*_ versus *F*_*Z*_ during contact are similar to those during allopatry in Fig. 5. However, *I*_*Z*_ values are typically larger than those for butterflies (Fig. 1) at intermediate values of *F*_*Z*_; Results for *s* = 0.1 and Δ*t*_*A*_ = 2*N* are shown in Fig. 8 (note that data is recorded in the gap region, *F*_*Z*_ *≤* 0.3, consistent with Fig. 6). As expected [11], larger rates of crossing over between genes, consistent with larger distances between genes on butterfly chromosomes, lead to smaller differences, Δ*F*, between *F*_*Z*_ and *F*_*A*_ (Fig. 9). Interestingly, the “null model” leads to smaller values of Δ*F* during contact (Fig. 9) than allopatry (Fig. 5). Lower rates of migration, for which populations tend toward continued speciation, are briefly explored in Fig.s S6–S7. In this case, populations do not exhibit “fat tail” statistics for low values of *F*_*A*_, and large values of Δ*F* are less frequent.

**Fig. 6.**
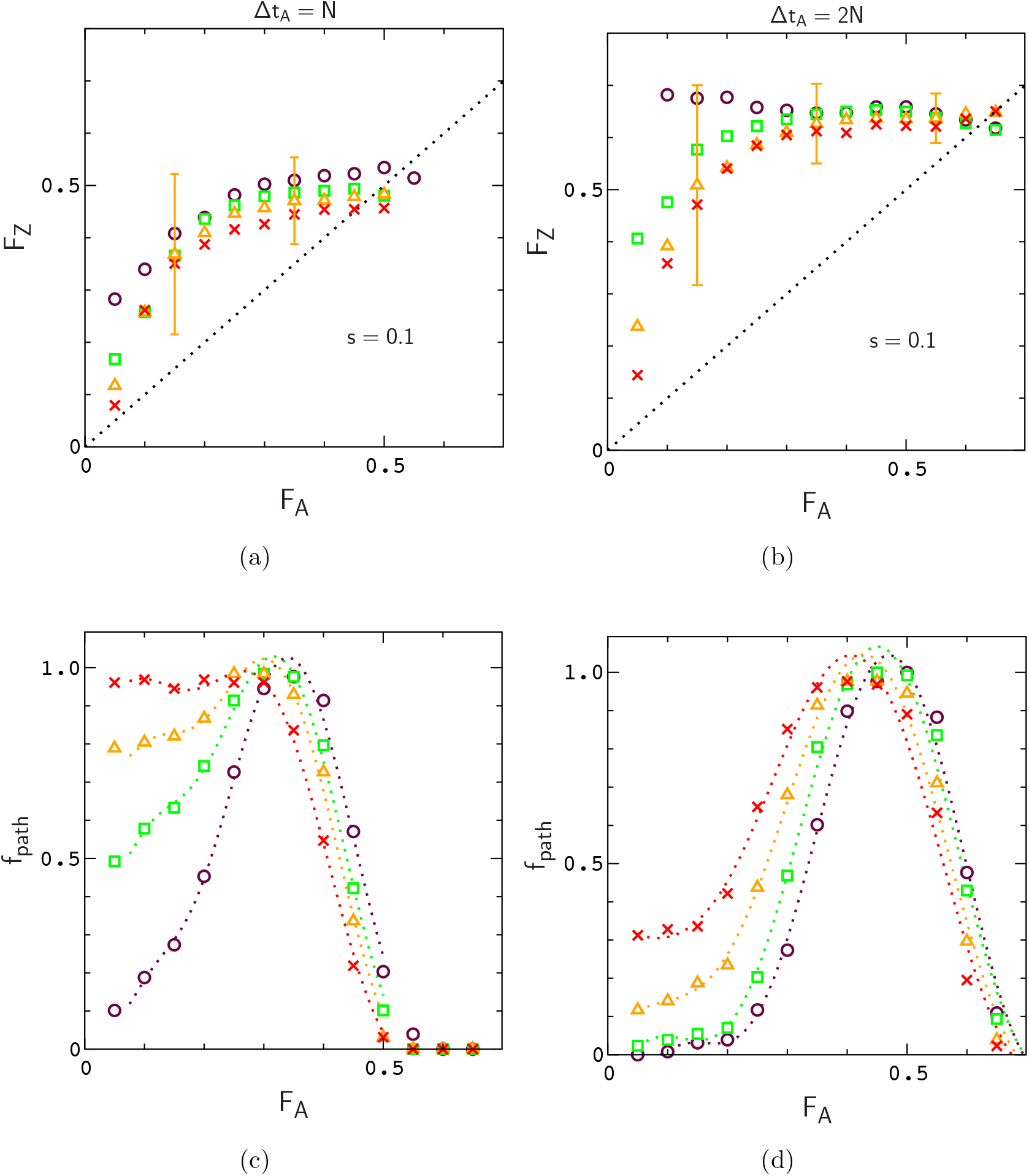
Mean value of *F*_*Z*_ versus *F*_*A*_ for several values of the migration rate, *N_ϵ_* = 1.5 (circles), 2.5 (squares), 4 (triangles), and 6 (crosses), and different times in allopatry. Each set of averages is computed from 128 replicate simulations with *N* = 10^4^, *µ* = 10^*−*4^, *r, r*^*′*^ = 10^*−*2^ and Δ*t*_*C*_ = *N*. Error bars describe the widths of the distributions for *NE* = 4. Lower panels describe the fraction of simulation paths contributing to each bin for *F*_*A*_.

**Fig. 7.**
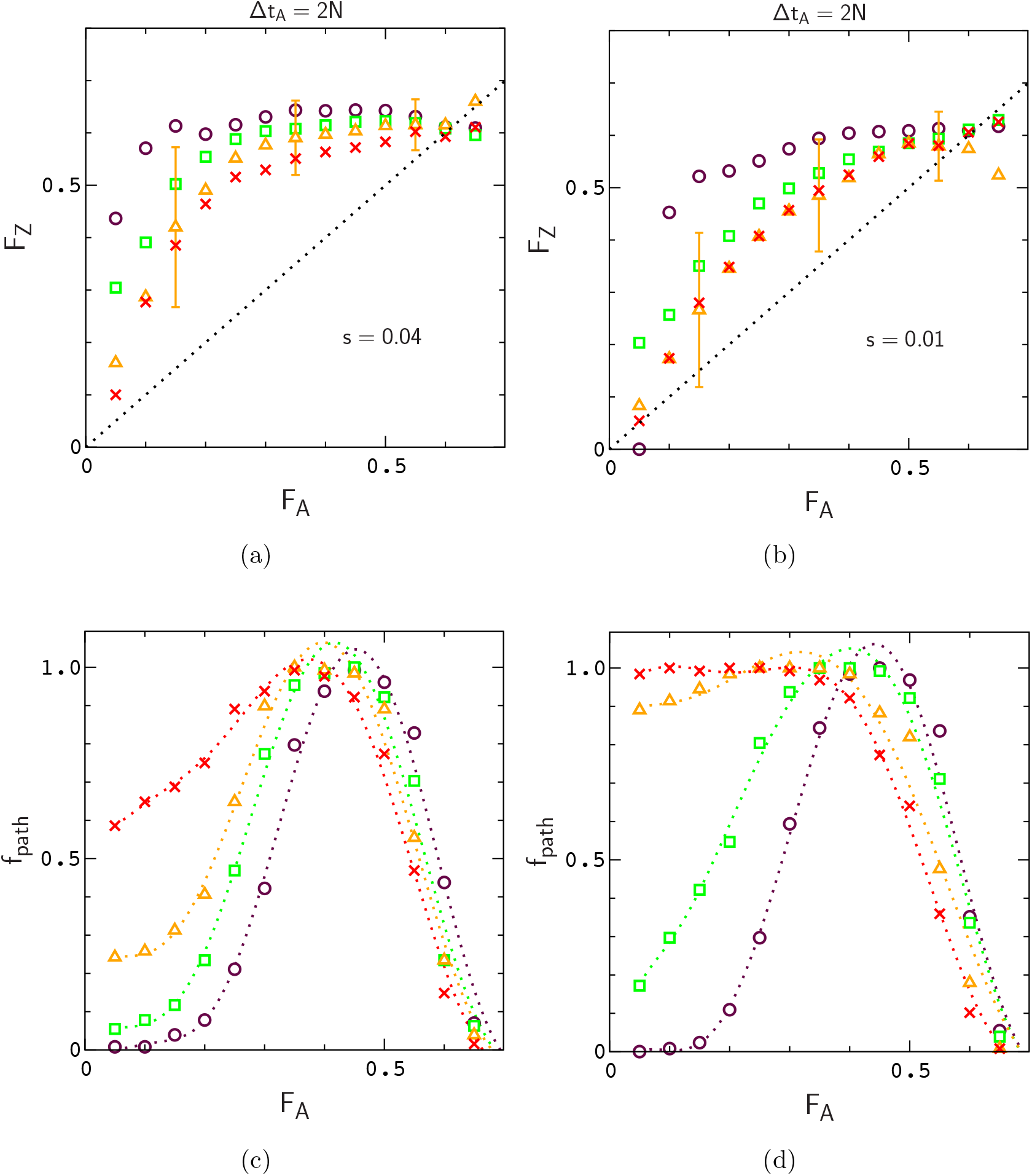
Mean value of *F*_*Z*_ versus *F*_*A*_ for decreasing interaction strengths. The parameters of the simulations are the same as those listed in Fig. 6 except where indicated in the figure.

**Fig. 8.**
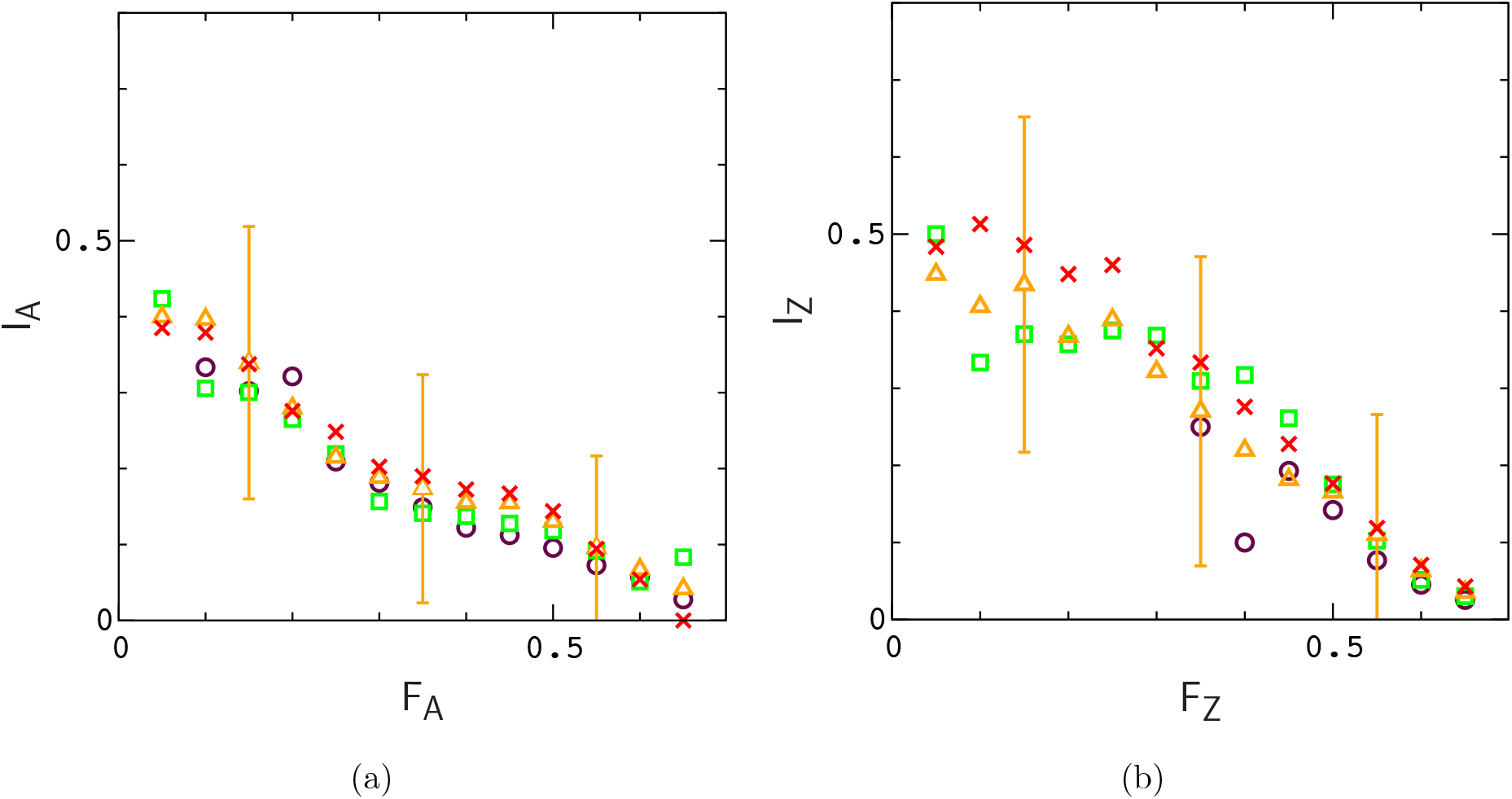
Mean values of *I*_*A*_ versus *F*_*A*_ and *I*_*Z*_ versus *F*_*Z*_ for the simulations in Fig. 6B. Error bars indicate the widths of the distributions for *Nϵ* = 4. The widths reflect the relatively small numbers of gene sequences used to determine *I*_*A*_ and *I*_*Z*_.

**Fig. 9.**
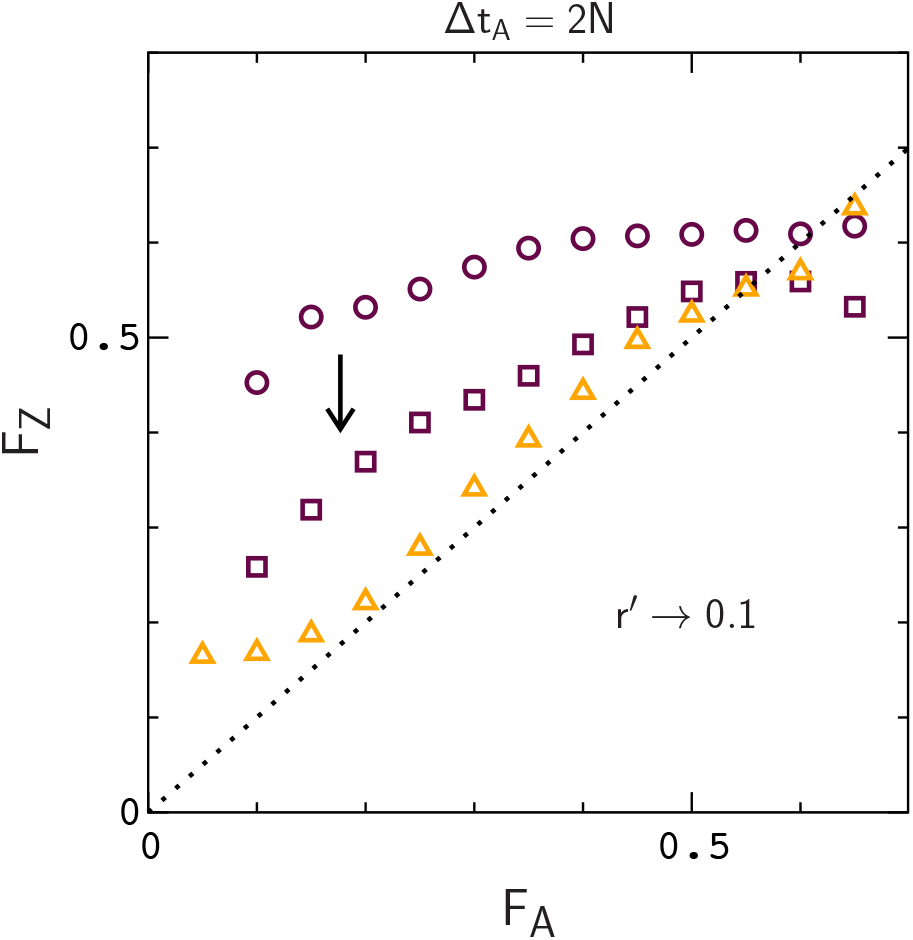
Mean value of *F*_*Z*_ versus *F*_*A*_ as the rate of crossing over between gene sequences is increased from *r*^*′*^ = 0.01 (circles) to *r*^*′*^ = 0.1 (squares). Averages are computed from 128 replicate simulations with *s* = 0.01, *N* = 10^4^, *µ* = 10^*−*4^, *r* = 10^*−*2^, and Δ*t*_*C*_ = *N*. Results for the “null model” (*s* = 0) for *r*^*′*^ = 0.01 (triangles) are included for comparison. Additional statistics are provided in Fig. S5.

In planning our simulations, we have implicitly assumed that real populations are subjected to repeated periods of isolation and contact on the order of glacial and interglacial periods. Given that real populations may be in various stages of divergence, we studied a single cycle of isolation and contact as a means to determine the influence of large–Z hybrid interactions during periods of contact. Because the morphologies of butterfly sister species (i.e., sex organs, wing color patterns, etc.) are similar, species boundaries are presumed to be somewhat porous, consistent with higher rates of migration in the model. For sufficiently strong interactions, the model leads to large differences between *F*_*Z*_ and *F*_*A*_ under fusion conditions, comparable, on average, to those in Fig. 2. In addition, the gap region, *F*_*Z*_ *≤* 0.3 in Fig. 1 is statistically unlikely in the model over a wide range of interaction strengths and migration rates. However, while mean values of *I*_*A*_ during contact are consistent with the data for *I*_*A*_ in Fig. 1A, the results for *I*_*Z*_ are clearly different from Fig. 1B for intermediate values of *F*_*Z*_, and appear to reflect a missing feature, or features, in the model dynamics. For example, if male butterflies are more abundant than females, effective population sizes and levels of introgression on autosomes and Z chromosomes will tend be similar, which should lead to smaller values of *I*_*Z*_. Increasing the numbers of genes segments considered in the model may also lead to better results. In any case, the model is clearly a very simplistic representation of butterfly dynamics.

In real populations [2] butterflies forage and interbreed on large structured landscapes, often consisting of connected islands on which butterfly numbers can vary dramatically (see, for example [20–22]). During periods of isolation, sister populations may evolve discordant mating cycles [13] and mate preferences [1], which limit interbreeding during periods of contact. Organisms can also evolve preferences for local resource types [23], which can limit migration across contact zones, or cause asymmetry in migration rates. Although some of these effects have been explored using island, or stepping stone models [10, 11, 24, 25], it would be interesting to know how interbreeding populations evolve on continuous landscapes, including basic behavioral aspects of butterflies (e.g. flight patterns [21], mating cycles, etc.), and the topographies of their resource distributions. The present work is a first step toward this goal.

## Acknowledgments

It is a pleasure to thank Jing Zhang and two anonymous reviewers for helpful comments during the completion of this work. This study is supported in part by a grant (to NVG) from the National Institutes of Health (GM127390).

