## supplemental figures for "Influence of the large–Z effect during contact between butterfly sister species"

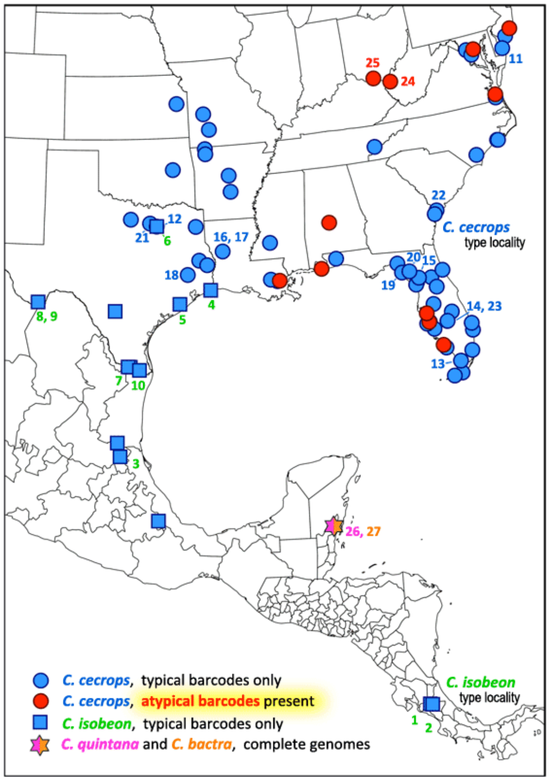


**Fig. S1.** Sample locations for Calycopis isobeon and Calycopis cecrops, the most frequently sampled pair of sister species studied by Cong et al. [1, 2]

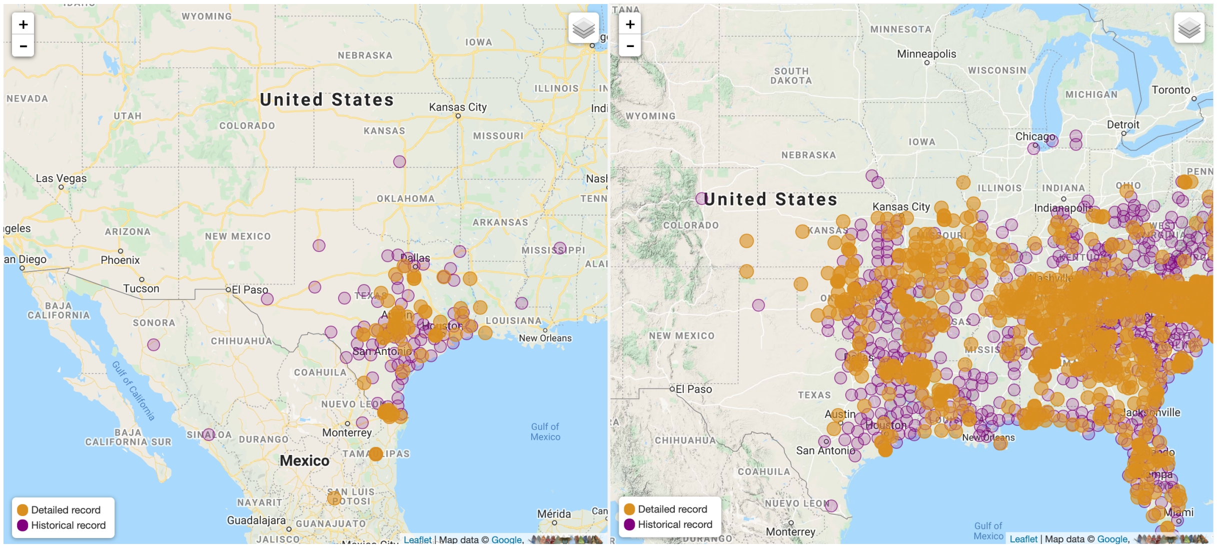


**Fig. S2.** Curated record of C. isobeon (left) and C. cecrops (right) butterfly sightings (source: Butterflies and Moths of North America, https://www.butterfliesandmoths.org).

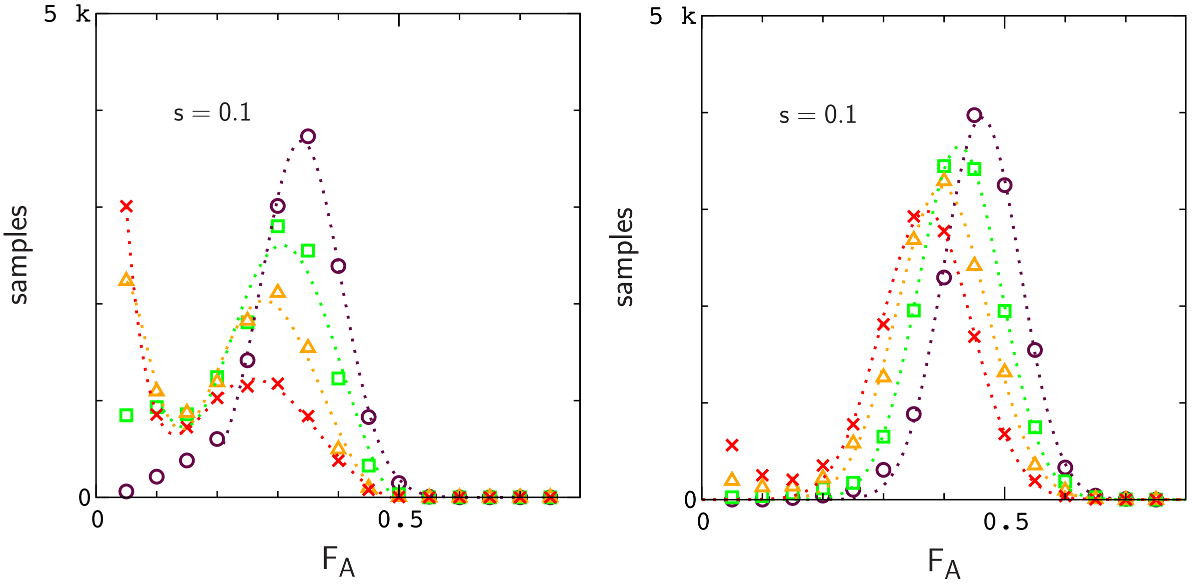


**Fig. S3.** Histograms of samples corresponding to Fig.s 6A (left) and 6B (right).

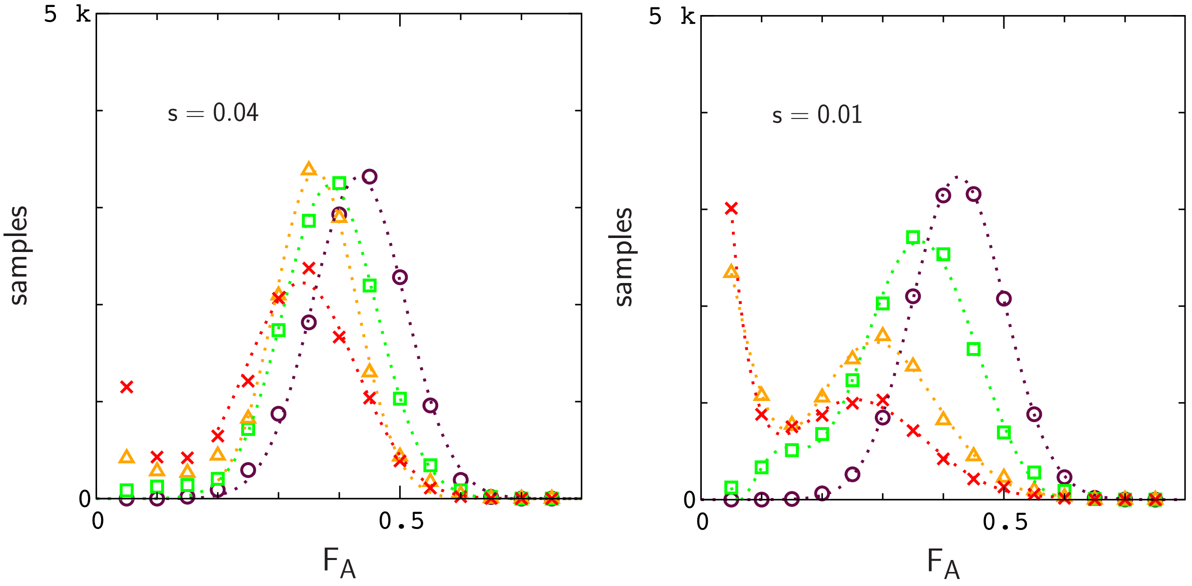


**Fig. S4.** Histograms of samples corresponding Fig.s 7A (left) and 7B (right).

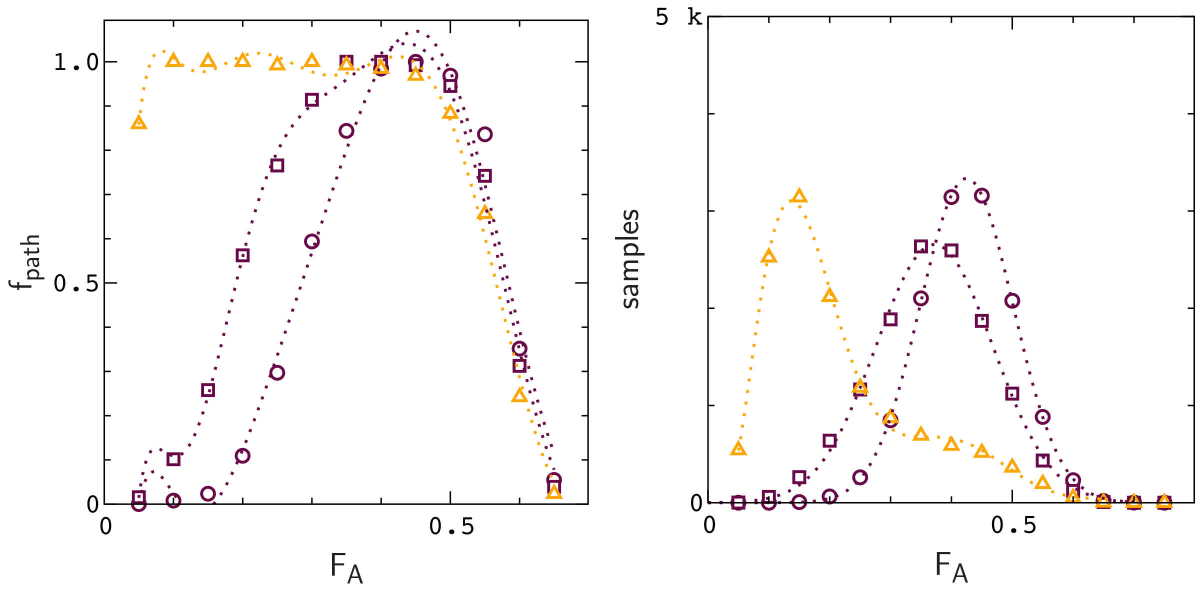


**Fig. S5.** Fraction of simulation paths (left) and histogram of samples (right) for Fig. 9.

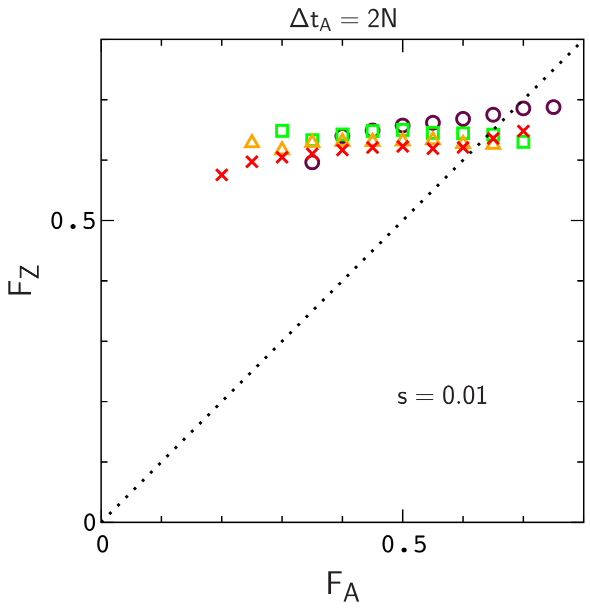


**Fig. S6.** Mean value of $F_Z$ versus $F_A$ for weak migration rates, $N\epsilon = 0.125$ (circles), 0.25 (squares), 0.5 (triangles), and 0.75 (crosses). Each set of averages is computed from 128 replicate simulations with $N = 10^4$, $\mu = 10^{-4}$, $r$, $r' = 10^{-2}$ and $\Delta t_C = N$.

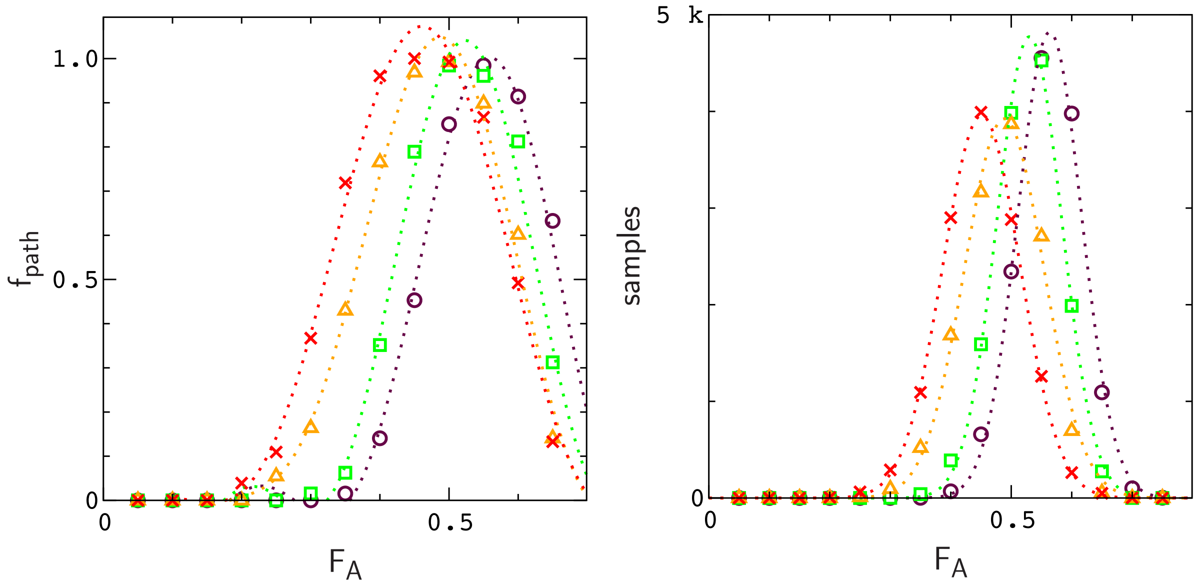


**Fig. S7.** Fraction of simulation paths (left) and histogram of samples (right) for Fig. S6.
